# Ecological and evolutionary patterns in the enigmatic protist genus *Percolomonas* (Heterolobosea; Discoba) from diverse habitats

**DOI:** 10.1101/612531

**Authors:** Denis V. Tikhonenkov, Soo Hwan Jhin, Yana Eglit, Kai Miller, Andrey Plotnikov, Alastair G.B. Simpson, Jong Soo Park

## Abstract

The heterotrophic flagellate *Percolomonas cosmopolitus* (Heterolobosea) is often observed in saline habitats worldwide that range from coastal waters to saturated brines. However, only two cultures assigned to this morphospecies have been examined using molecular methods, and their 18S rRNA gene sequences are extremely different. Further the salinity tolerances of individual strains are unknown. Thus, our knowledge on the autecology and diversity in this morphospecies is deficient. Here, we report 18S rRNA gene data on seven strains similar to *P. cosmopolitus* from seven geographically remote locations (New Zealand, Kenya, Korea, Poland, Russia, Spain, and the USA) with sample salinities ranging from 4‰ to 280‰, and compare morphology and salinity tolerance of the nine available strains. *Percolomonas cosmopolitus*-like strains show few-to-no consistent morphological differences, and form six clades separated by often extremely large 18S rDNA divergences (up to 42.4%). Some strains grew best at salinities from 75 to 125‰ and represent halophiles. All but one of these belonged to two geographically heterogeneous clusters that formed a robust monophyletic group in phylogenetic trees; this likely represents an ecologically specialized subclade of halophiles. Our results suggest that *P. cosmopolitus* is a cluster of several cryptic species (at least), which are unlikely to be distinguished by geography. Interestingly, the 9 *Percolomonas* strains formed a clade in 18S rDNA phylogenies, unlike most previous analyses based on two sequences.

## Introduction

The modern classification of eukaryotic microbes is usually based on molecular sequence information combined with morphological characters obtained using diverse imaging techniques, often including electron microscopy. Many taxonomists have a fundamental question whether small morphological differences among the observed organisms represent multiple species or one variable species [1–3]. Conversely, in protistology, many species (and genera) that were originally proposed based on light microscopy alone have proved to encompass considerable genetic diversity. Also, many morphospecies of protists are cosmopolitan and/or are found across a very wide range of habitats, raising the possibility of recognizing species that are divided more by geography or ecology than morphology [4]. Bickford et al. [1] regarded ‘cryptic species’ as cases where two or more species are distinguished that were previously assigned to a single morphologically defined species (i.e. morphospecies). The concept of cryptic (usually genetically different) species has been examined in a diverse range of protists, e.g. [4, 5, 6]. This concept is widely accepted due to an inconsistency between morphospecies and their DNA sequencing data. Obviously, the estimated number of protist species will vary tremendously depending on the species concept employed [7, 8].

The species *Percolomonas cosmopolitus* (formerly *Tetramitus cosmopolitus* Ruinen [9]) is a heteroloboseid flagellate with one long and three shorter flagella at the head of a ventral feeding groove, and no known amoeba stage in its lifecycle. Cell size is given as 6–12 μm long and 3–9 μm wide in the seminal modern accounts [10, 11]; the largest cells reported by Ruinen [9] are somewhat longer. This species is a particularly interesting protist for taxonomists and ecologists for several reasons: Firstly, it is possible that *P. cosmopolitus sensu lato* may be a remarkably broad assemblage of cryptic species. To date, two strains identified as *P. cosmopolitus* have been studied using molecular methods, but their 18S rRNA gene sequences share remarkably low sequence identity (58.4%). Secondly, *P. cosmopolitus* has been reported in samples from an extremely wide range of saline habitats, from coastal waters to saturated brines, throughout the world [9–12]. It is unknown whether there are ecologically or geographically defined subtypes. Thirdly, in addition to the genetic divergence between them (see above) *P. cosmopolitus* strains are usually not inferred to be sisters in phylogenetic trees, forming instead a paraphyletic group with respect to the pseudociliate *Stephanopogon*, which has many more flagella and a completely different feeding system [13–21]. Consequently, *Percolomonas cosmopolitus* and similar organisms (e.g. [22]) are interesting candidates to study the interplay between species distinctions, autecology and evolutionary history among protists.

Here, we examined seven *Percolomonas cosmopolitus*-like organisms, which were isolated from samples with a salinity range from 4‰ to 280‰, from seven different countries (New Zealand, Kenya, Korea, Poland, Russia, Spain, and the USA). The nine *Percolomonas* strains examined (including the two *P. cosmopolitus* strains previously sequenced) were morphologically similar to each other, but their 18S rRNA gene sequences showed considerable genetic divergence, and they formed six genetically distinct clusters. The strains isolated from high salinity samples preferred to grow in artificial media with salinity higher than seawater (75‰ or above), indicating ecological specialization. *Percolomonas* formed a monophyletic group in the 18S rRNA gene phylogeny.

## Materials and methods

### Isolation and cultivation

Seven monoprotistan strains were isolated from brackish to high salinity (4–280‰) water/sediment interface samples collected from New Zealand, Kenya, Korea, Poland, Russia, Spain, and the USA between 2008 and 2015 (Table 1). Isolates from higher salinity waters were LRS, SD2A, XLG1-P, S4, and P5-P; isolates from lower salinity waters were LO, and HLM-6. Strain HLM-6 was examined morphologically by A.P. Mylnikov [22] under the name *Percolomonas lacustris*. Each monoprotistan culture was established by single-cell isolation (from raw samples or crude cultures) or by serial dilution. High salinity media (~100‰ salinity) was made by dilution of Medium V (300‰; 272g NaCl, 7.6g KCl, 17.8g MgCl_2_, 1.8g MgSO_4_·7H_2_O, 1.3g CaCl_2_ l^−1^ water, see [23]) with sterile distilled water, whereas normal salinity media (~35‰ salinity) used autoclaved seawater. Luria-Bertani Broth (final concentration of 0.5%, Difco) plus autoclaved barley grains were added into each media to grow indigenous prokaryotes in the culture. For maintenance of each culture, 0.1 ml of inoculum was added into 5 ml of 100‰ salinity liquid media (isolates LRS, SD2A, XLG1-P, S4, and P5-P) or 35‰ salinity liquid media (isolates LO and HLM-6 plus the already available strains ATCC 50343 and White Sea = ‘WS’). Strain HLM-6 was also maintained in marine Schmalz-Pratt’s medium (NaCl–28.15 g l^−1^, KCl–0.67 g l^−1^, MgCl_2_·6H_2_O–5.51 g l^−1^, MgSO_4_·7H_2_O–6.92 g l^−1^, CaCl_2_·H2O–1.45 g l^−1^, KNO_3_–0.1 g l^−1^, K_2_HPO_4_·3H_2_O–0.01 g l^−1^ with a final salinity of 35‰) with addition of *Pseudomonas fluorescens* bacteria as food. All cultures were incubated at 25 °C and subcultured every four weeks.

**Table 1.**
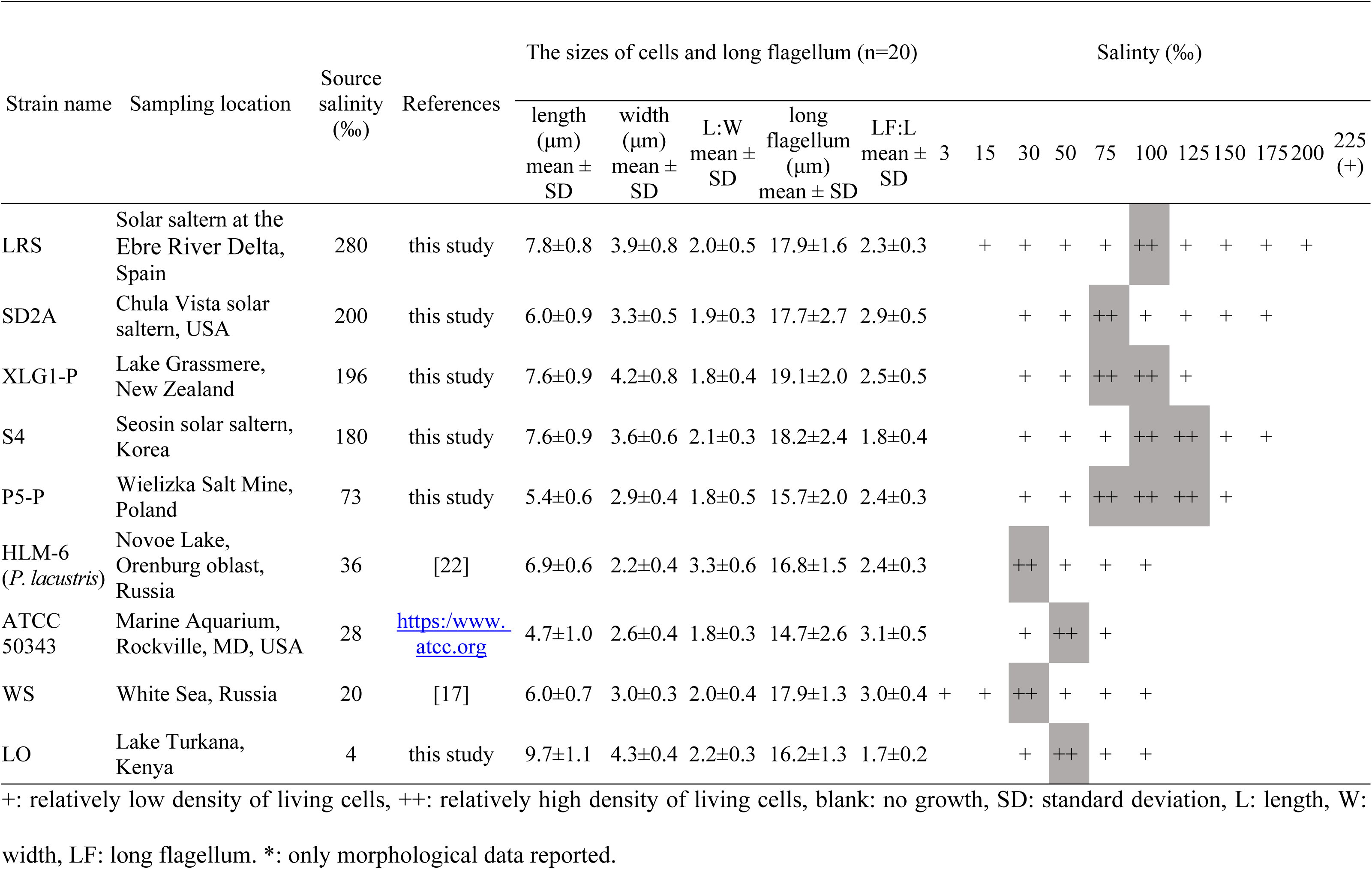
Characterizations of *Percolomonas* strains; sizes and salinity range supporting growth.

### Light microscopy

Live flagellates mounted on glass slides were observed with phase contrast microscopy or differential interference microscopy using a Leica DM5500B microscope equipped with a DFC550 digital camera (Leica, Wetzlar, Germany) or Carl Zeiss AxioScope A.1 microscope equipped with a AVT HORN MC-1009/S analog video camera. To observe the number and shape of the flagella, cultures (1 ml) were centrifuged at ×2,000 *g* for 10 min, then 900 µL of the supernatant was discarded, and the remaining volumes (i.e. 100 µL) were fixed by addition of 50 µL of 25% v/v glutaraldehyde (electron microscopy grade). The sizes of the live cells (i.e. 20 cells per culture) were measured from digital images.

### Scanning electron microscopy

Cultures (1 ml) were centrifuged at ×1,200 *g* for 10 min, then 900 µL of the supernatant was discarded, and the remaining volumes (i.e. 100 µL) were fixed by adding 50 µL of 25% v/v glutaraldehyde (electron microscopy grade). Fixed cells were allowed to settle (40 min) on glass coverslips coated with 1% poly-L-lysine. Cells were rinsed with sterile media, and then dehydrated with a graded ethanol series (30–100%). The glass coverslips were then critical-point dried. Fixed cells were coated with gold/platinum using an ion sputter system. Specimens were examined with a SU8220 field emission scanning electron microscope (Hitachi, Tokyo, Japan) or JSM-6510LV scanning electron microscope (JEOL Ltd., Tokyo, Japan).

### Molecular sequencing and phylogenetic analysis

Nucleic acids from the seven new isolates were extracted using a DNeasy Blood and Tissue Kit (Qiagen, Hilden, Germany) or MasterPure Complete DNA and RNA Purification Kit (Epicentre, Madison, USA), as described in the supplied protocols.

For all strains except LRS and HLM-6, the 18S rRNA gene sequences were obtained by PCR amplification using a combination of the eukaryote primers EukA 5′-AACCTGGTTGATCCTGCCAGT-3′ and EukB 5′-TGATCCTTCTGCAGGTTCACCTAC-3′ [24]. The 20-µL PCR reactions included 1.5 µL each of 10-µM stocks of the primers, 2 µL of a 0.25-mM dNTP-mix, 0.8 µL of 50 mM MgCl_2_, 0.2 µL of 5 U/µL *Taq* DNA polymerase (Solgent, Daegeon, Republic of Korea), and 1–3 µL of DNA template. The cycling conditions were as follows: an initial denaturing step at 94 °C for 5 min, followed by 35 cycles of 30 s at 94 °C, 1 min of annealing at 55 °C, and extension at 72 °C for 2 min, with a final extension step for 10 min at 72 °C. Amplicons were cloned into a pGEM-T Easy vector, at least five positive clones per sample were partially sequenced, and a positive clone was completely sequenced using various sequencing primers. For strain XLG1-P, the cycling conditions were slightly different: 35 cycles of 20 s at 94 °C, 1 min of annealing at 55 °C, and 3 min of extension at 72 °C. Strain LRS was amplified using the different eukaryote primers 82F (5’-GAAACTGCGAATGGCTC-3’) and 1498R (5’-CACCTACGGAAACCTTGTTA-3’). The optimized PCR condition was 2 min at 96 °C (an initial denaturation), followed by 35 cycles of 30 s at 96 °C, 1 min at 60 °C, 2 min at 72 °C, with a final extension for 10 min at 72 °C. The PCR products for strains XLG1-P and LRS were directly sequenced by Sanger dideoxy sequencing without cloning.

For Strain HLM-6, the primers used were PF1 5′-GCGCTACCTGGTTGATCCTGCC-3′ and FAD4 5′-TGATCCTTCTGCAGGTTCACCTAC-3′ [25, 26]. The 25-µL PCR reaction included 0.5 µL each of 10-µM primer stocks, 1 µL of DNA template, 10.5 µL PCR-grade water and 12.5 µL EconoTaq® PLUS Green 2x Master Mix (Lucigen, Middleton, USA). The cycling conditions were 95 °C for 3 min, followed by 35 cycles of 30 s at 95 °C, 30 s at 50 °C, and 1.5 min at 72 °C for 1.5 min, with a final extension of 5 min at 72 °C. Amplicons were cloned using Strata Clone PCR Cloning Kit (Agilent, Santa Clara, USA). The 18S rRNA gene sequences from the isolates have been deposited in GenBank under the accession numbers XXXXXXX.

The 18S rRNA gene sequences from 62 representative heterolobosean species, plus 16 other Discoba species selected as outgroups, were used for phylogenetic analysis (the seed alignment originated from Jhin and Park [16]). The dataset was aligned and masked by eye, with 1,033 unambiguously aligned sites retained for analysis. The alignment is available on request. Phylogenetic trees were inferred by Maximum Likelihood (ML) and Bayesian analyses. The GTR + gamma + I model of sequence evolution was selected for the dataset using MrModeltest 2.2 [27] and was used for both analyses. The ML tree was estimated using RAxML-VI-HPC v.7 [28] with the GTRGAMMAI model setting, 500 random starting taxon addition sequences, and statistical support assessed using bootstrapping with 10,000 replicates. The Bayesian analysis was conducted in MrBAYES 3.2 [29] with two independent runs, each with four chains running for 2 × 10^7^ generations with the default heating parameter (0.1) and sampling frequency (0.01). A burn-in of 30% was used, by which point convergence had been achieved (the average standard deviation of split frequencies for the last 75% of generations was < 0.05).

### Salinity ranges for growth

To estimate the salinity ranges supporting growth of the seven new isolates and two previously available isolates, we performed an experiment using media with 3‰ to 300‰ salinity, made from artificial seawater stock (Medium V; see above) as reported by Park and Simpson [18, 30]. In brief, the medium was supplemented with heat-killed *Enterobacter aerogenes* at an initial density of 3.46 × 10^7^ cells per ml (20 µL) at 7- to 14-day intervals to support the growth of the protists. All treatments were performed in duplicate. Medium V (0.96 ml) with a range of salinities (3‰–300‰) were inoculated with 20 µL of actively growing stock culture (100‰ or 35 ‰ salinity media with autoclaved barley grain) and incubated in the dark at 25 °C for at least 49 days. We confirmed the salinity range supporting growth by transferring a sample of the isolate into fresh media with the same salinity (0.96 ml of media; inoculum size 20 µL), and re-examining the culture for actively moving cells at 7- to 14-day intervals over a period of 35 days.

## Results

### General morphology

Live cells were usually ovoid-shaped or spindle-shaped with average lengths and widths ranging from 4.7–9.7 µm and 2.2–4.3 µm (Fig 1 and Table 1). The ratio of length and width of the cells was between 1.8 and 3.3 on average (Table 1). The biggest cells (length: 9.7 µm, width: 4.3 µm on average) were from strain LO isolated from Lake Turkana, Kenya, whereas the smallest cells were from strain ATCC 50343 (length: 4.7 µm, width: 2.6 µm on average), but the sizes of different strains represented an overlapping continuum. Cells had four flagella inserted in the sub-anterior part of the cell, at the head of the ventral cytostomal groove (Fig 2). Three flagella were shorter, and one flagellum was longer. The three short flagella were similar in length, similar to that of the cytostomal groove (typically ~4 µm long). The long flagellum averaged 14.7–19.1 µm in length, depending on the strain, which was 1.7–3.1 times longer than the length of the cell body (Table 1). Most cells had an acroneme at the tip of the long flagellum (Fig 2), however, this seems not to be a permanent feature. All species showed jerking motility and sometimes rotated in a counterclockwise direction. The right side of the cytostomal groove was curved, whereas the left side was roughly linear (Fig 2). No amoeboid form was observed during the cultivation of the organisms, and a cyst form was observed occasionally in all cultures except for strain ATCC 50343. We did not observe discrete features that distinguished isolates from each other under a light microscope or a scanning electron microscope.

**Fig 1.**
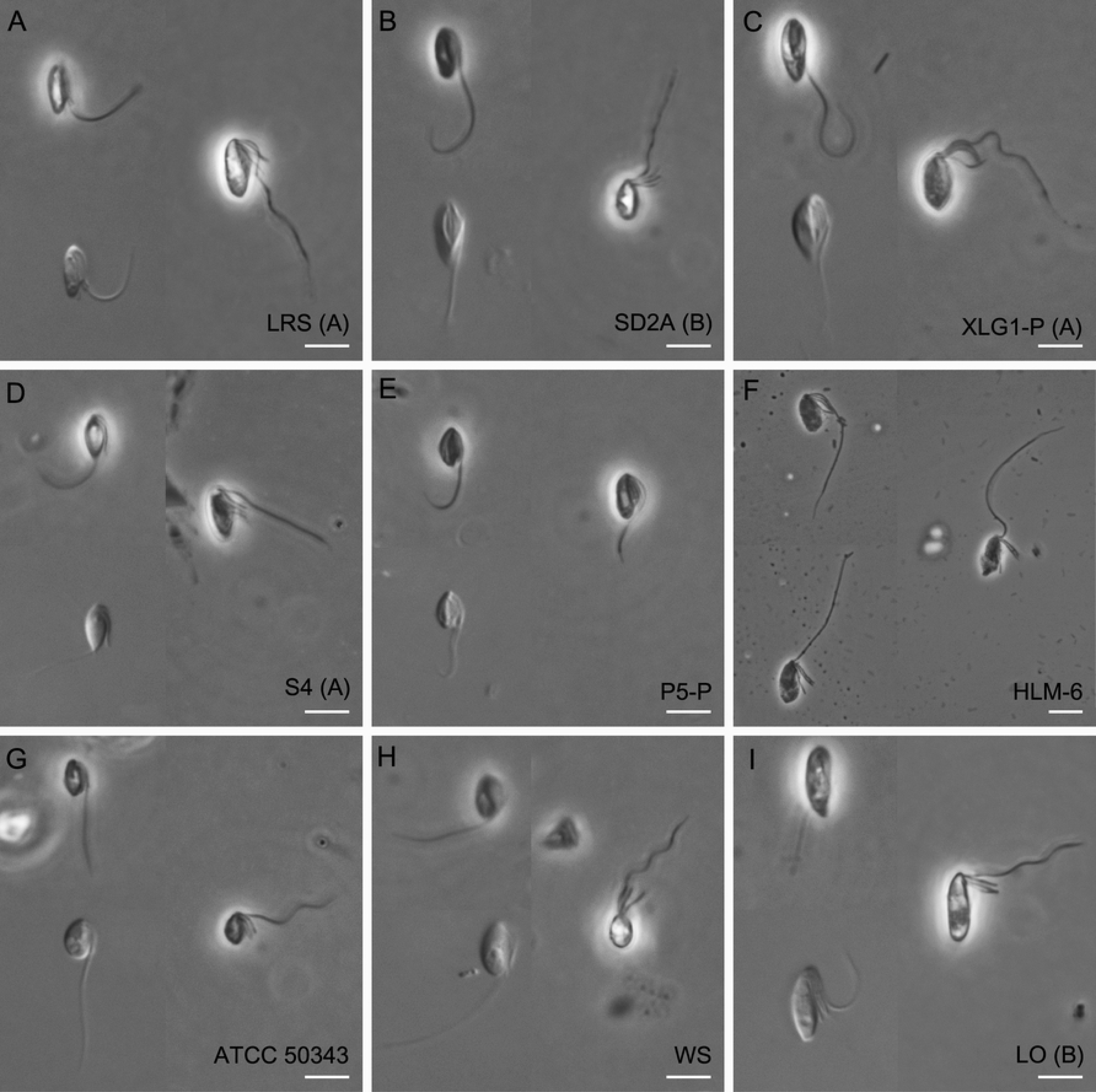
General light micrographs of *Percolomonas cosmopolitus*-like strains (ordered by source salinity as in Table 1). (A) New strain LRS isolated from a solar saltern at the Ebre River Delta, Catalonia, Spain (280‰ salinity). (B) New strain SD2A isolated from Chula Vista solar saltern, USA (200‰ salinity). (C) New strain XLG1-P isolated from Lake Grassmere, New Zealand (196‰ salinity). (D) New strain S4 isolated from a Korean solar saltern (180‰ salinity). (E) New strain P5-P isolated from Wielizka salt mine, Poland (73‰ salinity). (F) Strain HLM-6 isolated from Novoe Lake, Orenburg oblast, Russia (36‰ salinity); this strain was the type for *Percolomonas lacustris* as described by A.P. Mylnikov [22]. (G) “*Percolomonas cosmopolitus”* ATCC 50343. (H) “*Percolomonas cosmopolitus”* strain WS (White Sea), originally reported by Nikolaev et al. [17]. (I) New strain LO isolated from Lake Turkana, Kenya (4‰ salinity). Note that the right cell image in each panel represents a fixed cell. Groups (A) and (B) indicate halophile clades ‘A’ and ‘B’ in Fig. 3. All scale bars represent 5 μm.

**Fig 2.**
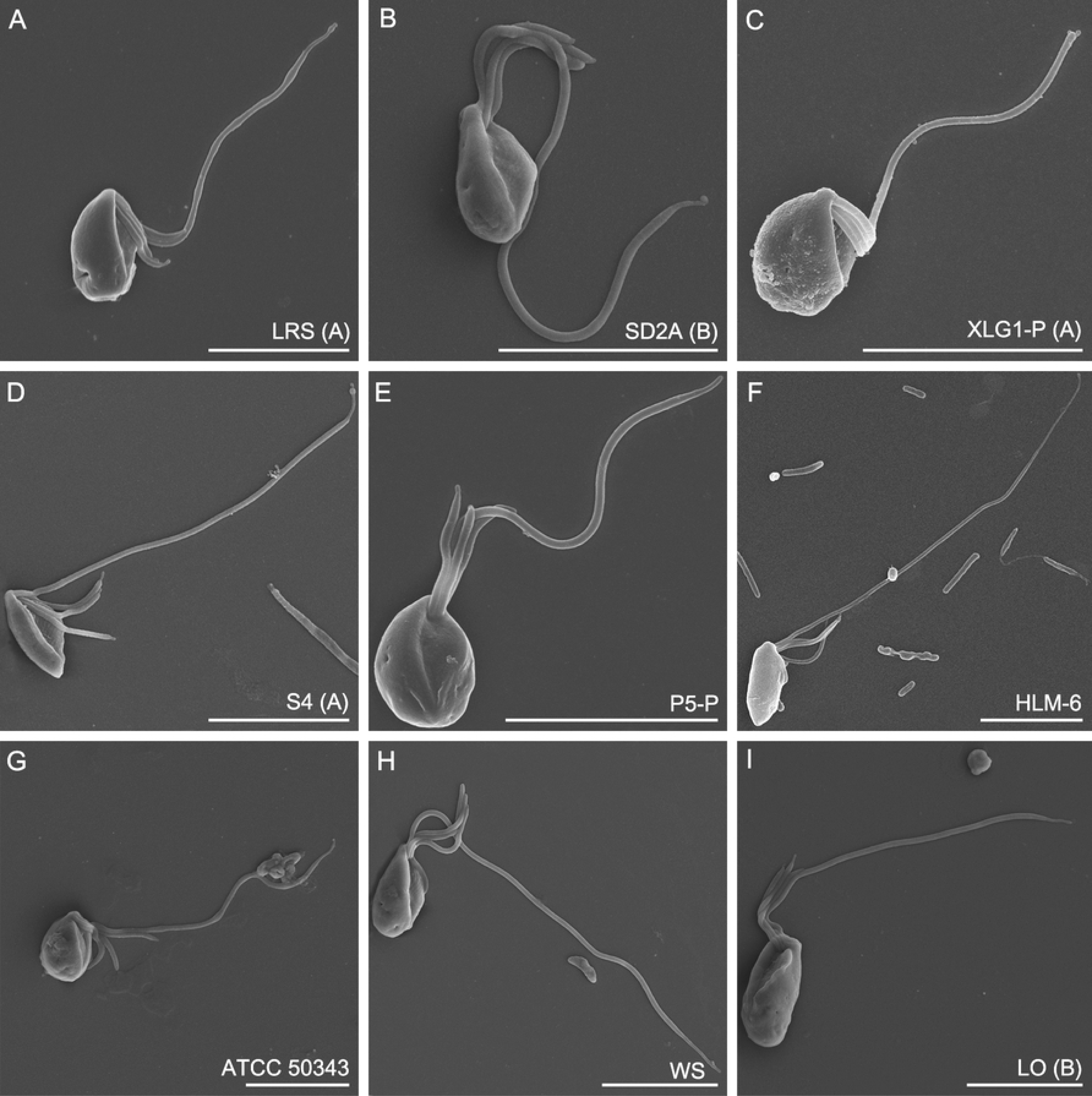
Scanning electron micrographs of *Percolomonas cosmopolitus*-like strains. (A) Strain LRS (B) Strain SD2A. (C) Strain XLG1-P. (D) Strain S4. (E) Strain P5-P. (F) Strain HLM-6 = *Percolomonas lacustris*. (G) ATCC 50343. (H) Strain WS. (I) Strain LO. The order of strains in Fig. 2 is the same as that in Fig. 1 and Table 1. All scale bars represent 5 μm.

### Molecular phylogeny of 18S rRNA gene sequences

The 18S rRNA gene sequences from the seven unstudied *Percolomonas* strains were closest by BLASTN search to *Percolomonas cosmopolitus* strain WS (AF 519443), but with a low identity of 74% to 83%. As in previous studies [14–16, 19, 31, 32], Heterolobosea formed a strong monophyletic group in phylogenetic trees of 18S rRNA gene sequences (Fig 3). The seven new *Percolomonas* strains and two previously identified sequences branched with the pseudociliate taxon Stephanopogonidae with strong support (100% ML; PP 1; Fig 3) forming the clade Percolatea. Interestingly, Percolomonadidae, including the seven new sequences, formed a monophyletic group, with moderate bootstrap support (ML: 76%) and posterior probability 1. Within Percolomonadidae, ATCC 50343 (USA) and P5-P (Poland) were distinct from each other and the remaining seven strains, which formed a maximally supported clade. This subdivided further into (i) WS (Russia), (ii) HLM-6 (Russia), and (iii) a maximally supported ‘putative halophile clade’ of 5 strains in 2 clusters, ‘A’ and ‘B’, for a grand total of 6 distinct clusters or single-strain lineages. The 18S rDNA sequences differences between these 6 clusters range from 15.1 to 42.4%. Group A of the putative halophile clade was composed of strains LRS (Spain), XLG1-P (New Zealand), and S4 (Korea) and formed a maximally supported clade (Fig 3), with low genetic divergence between the strain (98% to 99% identities). All members were isolated from hypersaline waters of 180 to 280‰ salinity (Table 1). Group B consisted of strains LO (Kenya) and SD2A (USA) which showed 98% sequence identity, though formed a weakly supported clade (62% ML; PP 0.75). Strain SD2A was isolated at 200‰ salinity, while strain LO was isolated from a low salinity (4‰) sample, though it cannot grow at this salinity (see below and Table 1).

**Fig 3.**
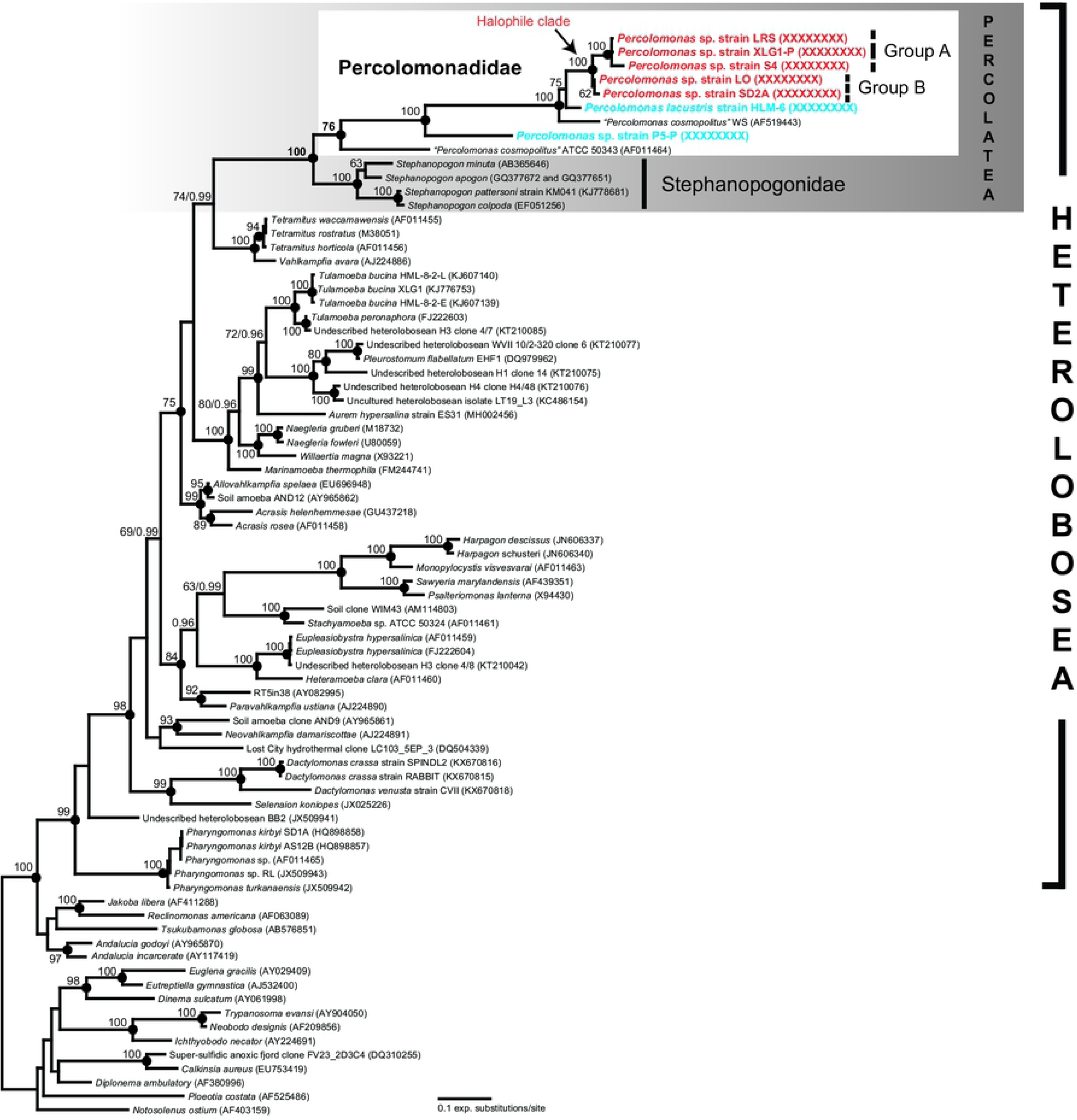
Maximum likelihood phylogenetic tree of 18S rRNA gene sequences from *Percolomonas* strains, other representative heterolobosean species, and outgroups (i.e. Euglenozoa, Jakobida, and *Tsukubamonas globosa*). Bootstrap support values (> 60%) are shown at the nodes. Solid circles represent a Bayesian posterior probability of 1 (posterior probability < 0.95 not shown). Note that ‘Group A’ and ‘Group B’ in Percolomonadidae together represent the putative halophilic *Percolomonas* clade.

### Salinity tolerance of *Percolomonas* strains

The *Percolomonas* strains were isolated from a variety of habitats with 4‰–280‰ salinity. Salinity ranges supporting growth of all nine sequenced *Percolomonas* isolates were determined as described earlier [16, 18, 20, 21]. *Percolomonas* strain LRS isolated from 280‰ salinity showed the broadest salinity range for growth (15‰–200‰ salinity, Table 1). Strain ATCC 50343 grew in the narrowest salinity range (30‰–75‰ salinity, Table 1). *Percolomonas* strain LO isolated from a source salinity of 4‰ grew only at 30‰ to 100‰ salinity. This suggests that the strain existed as an alternative life-cycle form (e.g. cyst) in the original sample. The five *Percolomonas* strains (i.e. LRS, SD2A, XLG1-P, S4, and P5-P) isolated from hypersaline habitats with >70‰ source salinity could grow best at 75‰ to 125‰ salinity (Table 1). In contrast, the four *Percolomonas* strains (i.e. ATCC 50343, White Sea, HML-6 and LO) isolated from non-hypersaline habitats with < 40‰ source salinity appeared to grow best at 30‰ to 50‰ salinity (Table 1).

## Discussion

### Morphology of *Percolomonas* isolates

In general, the isolates studied here are morphologically indistinguishable by light and scanning electron microscopy from the seminal modern accounts of *Percolomonas cosmopolitus* [10, 11]. They all have four flagella, specifically three short flagella (~4 μm long) and one long flagellum (12–20 μm long) directed along a ventral groove. The size of the cell body of *P. cosmopolitus* is 6–12 μm in length, 3–9 μm in width [10], overlapping with all of our isolates. The curved right side of the cytostomal groove differs from the left side with a near-linear shape. All strains move with a jerky gliding motion. Thus, it is appears that all new isolates may be assigned to the morphospecies *P. cosmopolitus*. The nominal *Percolomonas* species most similar to *P. cosmopolitus* include *P. denhami*, with three flagella, two of which are long [33] and *P. similis* with two flagella, one short, and the other long [34]. There are no molecular data for either of these species. The third similar species is *Percolomonas lacustris*, which was introduced based on the description of strain HLM-6. This was distinguished morphologically from *P. cosmopolitus* primarily by having a small posterior bulbous projection and an acroneme on one short flagellum [22]. However, the bulbous projection appears by light microscopy just as a pointed tip, which was a feature of the original description of *P. cosmopolitus* [9], and was seen intermittently in several of our isolates (see Fig 1). We confirmed that one short flagellum bearing a proportionately long acroneme can be a feature of strain HLM-6 (compare our Fig 2f to Fig 2g in [22]), but note that other cells of the strain appear to lack it (Fig 2f in [22]), and that (shorter) acronemes can be observed on the short flagella of several of the strains studied here (our Fig 2). Thus, it seems that the morphological distinction between *P. lacustris* and other *P. cosmopolitus*-like strains is subtle at best.

### Genetic structure in *Percolomonas*

The *Percolomonas* complex includes 6 distinct clades that are separated by genetic distances of 15.1% or more. Considering the similar morphology of *P. cosmopolitus*-like strains (see above), it is reasonable to regard the *Percolomonas* complex as a grouping of several (at least) cryptic (or sibling) species. Among protists, it has long been recognized that numerous cryptic species can exist within a single morphospecies [4, 8, 35–38]. Fenchel [36] suggested that the cryptic protists are regarded as allopatric speciation, and are distributed in limited geographic regions. In our case, however, no geographic clustering of *Percolomonas* strains was observed in the two clades represented by multiple isolates, although the numbers of sampling sites and sequencing data are limited. Three *Percolomonas* strains in ‘Group A’ form a robust clade with low genetic divergences (98% to 99% identities) and are widely dispersed geographically (Korea, New Zealand, and Spain). In addition, two *Percolomonas* strains in ‘Group B’ (with 98% identity) are from Africa and North America. This pattern is more consistent with extremely widespread dispersal, as inferred for other very small protists (e.g. [39]), and we speculate that environmental selection to different habitats may determine which groups appear where (see below).

### Halophilicity in *Percolomonas*

Ruinen [9] reported that *P. cosmopolitus* was detected at salinities ranging from 30‰ to saturated brines. In our study, the nine examined *Percolomonas* strains derived from a wide range of salinities, up to near-saturation. In general, the halophilicity of these strains was related to the salt concentration of their original habitats. Five *Percolomonas* strains (i.e. LRS, SD2A, XLG1-P, S4, and P5-P) were isolated from various habitats with a salinity range of 73‰ to 280‰. These all grew best at 75‰–125‰ by our qualitative estimation and most could still grow at 175‰ or even 200‰ on our experimental media. This clearly makes them halophiles according to the definition by Oren [40], in which a halophile could grow at 50‰ or higher and tolerated at 100‰ salinity. None were obligate halophiles, however, since all could also grow at 30‰ salinity. By contrast, three *Percolomonas* strains (i.e. HLM-6, ATCC 50343, and WS) isolated from non-hypersaline habitats (20‰ to 36‰ salinity) grew optimally at 30‰ or 50‰, and failed to grow above 100‰ salinity. Interestingly, *Percolmonas* strain LO grew optimally at 50‰ salinity, and grew up to 100‰ salinity, although it was isolated from a Lake Turkana, Kenya, with a very low salinity of 4‰. Phylogenetically, LO belongs within the putative ‘halophile clade’, and it is possible that it descended from a more halophilic ancestor (see below). Interestingly, the Lake Turkana sample was also the origin for *Pharyngomonas turkanensis*, a heterolobosean amoeba that grows best at 15–30‰ salinity, but is inferred to have descended from halophilic *Pharyngomonas* ancestor [41].

The taxon Heterolobosea includes a substantial proportion of the known halophilic or halotolerant eukaryotes [18–21, 41–44], and is interesting for examining the evolution of halophiles [16, 42, 43]. Recently, Kirby et al. [43] and Jhin and Park [16] suggested that the Tulamoebidae clade (*sensu lato*) in Heterolobosea were an example of a radiation of morphospecies that stemmed from a common halophilic ancestor. This clade included *Pleurostomum flabellatum, Tulamoeba peronaphora, Tulamoeba bucina*, and *Aurem hypersalina*, all with optimal salinities for growth of at least 150‰ [16, 20, 21, 43]. It is possible that the *Percolomonas* clade consisting of ‘Group A’ and ‘Group B’ may also represent a radiation of halophiles (albeit one of cryptic species within a morphospecies). All of the cultivated *Percolomonas* in this clade are halophiles, with one borderline case (LO; see above). However, this possibility of an exclusively/predominantly halophile clade could be a biased sampling artefact, and should be tested through additional isolations and study of related strains. This would also be useful to understand the nature of the closest relative of the halophile strain P5-P, which is phylogenetically isolated from others at present.

### Is *Percolomonas* monophyletic?

For a long time only two 18S rRNA gene sequences of *P. cosmopolitus* were available, and these were included in many phylogenetic analyses of Heterolobosea [13–21, 45]. Most of these phylogenies showed the two sequences forming a paraphyletic group, with one more closely related to *Stephanopogon* [45]. This inference may have been affected by the small number of sequences of *Percolomonas* available. In the present study, with seven additional and different 18S rRNA gene sequences, we instead inferred a (moderately supported) *Percolomonas* clade. Future research will address the cause of this difference, and test the relationships amongst *Percolomonas* and *Stephanopogon* using other markers.

## Conclusions

On the basis of light and scanning electron microscopic observations, all *Percolomonas* strains studied here are morphologically very similar, in spite of the huge genetic diversity they encompass. *Percolomonas* strains form at least 6 genetically distinct clades in the molecular phylogenetic trees, and could be considered to represent at least as many cryptic species. The speciation of *Percolomonas* strains could be partially related to salinity preference, rather than spatial distribution. The clusters ‘Group A’ and ‘Group B’, which are specifically related, may collectively represent a halophilic clade.

## Acknowledgements

We thank Dr. Alexander Mylnikov for help with SEM preparations and Dr. Jan Janouškovec for help with SSU rDNA sequencing of strain HLM-6. We thank Dr. Noèlia Carrasco for providing access to the Delta de l’Ebre sampling location, and Golara Sharafi for isolating and obtaining DNA from strain LRS. This research was supported by the Basic Science Research Programs through the National Research Foundation of Korea (NRF), a grant from the National Institute of Biological Resources (NIBR), funded by the Ministry of Education (2016R1A6A1A05011910, 2017K2A9A1A06049946, and 2019R1A2C2002379) and the Ministry of Environment (MOE) of the Republic of Korea (NIBR201801208), respectively, to J.S.P., a Russian Foundation for Basic Research grant (No 18-504-51028) to D.V.T., and a Natural Sciences and Engineering Research Council of Canada Discovery Grant (298366-2014) to A.G.B.S.

## Author contributions

Conceptualization: Jong Soo Park, Alastair G.B. Simpson

Investigation: Jong Soo Park, Denis V. Tikhonenkov, Soo Hwan Jhin, Yana Eglit, Kai Miller, Andrey Plotnikov

Formal analysis: Jong Soo Park, Denis V. Tikhonenkov, Soo Hwan Jhin

Visualization: Jong Soo Park, Denis V. Tikhonenkov, Soo Hwan Jhin

Supervision: Jong Soo Park, Alastair G.B. Simpson

Funding acquisition: Jong Soo Park, Denis V. Tikhonenkov, Alastair G.B. Simpson

Writing – original draft preparation: Jong Soo Park, Denis V. Tikhonenkov, Alastair G.B. Simpson

Writing – review and editing: all authors.

